# Egg-laying by female *Aedes aegypti* shapes the bacterial communities of breeding sites

**DOI:** 10.1101/2022.02.22.481482

**Authors:** Katherine D. Mosquera, Luis Eduardo Martínez Villegas, Gabriel Rocha Fernandes, Mariana Rocha David, Rafael Maciel-de-Freitas, Luciano A. Moreira, Marcelo G. Lorenzo

## Abstract

*Aedes aegypti*, the main vector of multiple arboviruses, is highly associated with human dwellings. Females exhibit an opportunistic oviposition behavior, seldomly laying eggs on natural containers, but rather distributing them among human-generated breeding sites. Bacterial communities associated with such sites, as well as the compositional shifts they undergo through the development of larval stages, have been described. Some bacteria can play a direct role in supporting the success of mosquito development. Additionally, exposure to different bacteria during larval phases can have an impact on life-history traits. Whether the larvae acquire symbionts from aquatic niches, or just require bacteria as food, is still debated. Based on these facts, we hypothesized that female *Ae. aegypti* shape the bacterial communities of breeding sites during oviposition as a form of niche construction to favor offspring fitness. Our study presents a series of experiments to address whether gravid females modify bacterial consortia present in larval habitats. For this, we first verified if females can mechanically transfer bacteria into culture media. As evidence of mechanical transmission was obtained, we then elaborated an experimental scheme to dissect effects from factors related to the act of oviposition and mosquito-egg-water interactions. The DNA samples obtained from breeding site water belonging to five treatments were subjected to amplicon-oriented sequencing to infer their bacterial community structure. Microbial ecology analyses revealed significant differences between treatments in terms of diversity. Particularly, between-treatment shifts in abundance profiles were detected, also showing that females induce a significant decrease in alpha diversity through oviposition. In addition, indicator species analysis pinpointed bacterial taxa with significant predicting values and fidelity coefficients for the samples in which single females laid eggs. Furthermore, we provide evidence regarding how one of these indicator taxa, *Elizabethkingia*, exerts a positive effect upon the development and fitness of mosquito larvae, thus suggesting that the developmental niche construction hypothesis may hold in this model.

## INTRODUCTION

The mosquito *Aedes aegypti* (Linnaeus, 1762) is the main vector of several arboviruses such as those causing dengue, yellow fever, Zika, and chikungunya. Because of its wide distribution across tropical and subtropical regions and close association with urban areas, this mosquito represents a major threat to the health of millions of people annually (WHO, 2017; Kamal et al., 2018). Human dwellings and their surroundings represent suitable mosquito habitats in which there is a reduced number of predators, diverse sugar sources can be reached, hosts are widely available for blood-feeding (as well as resting places for gravid females), and a variety of water-holding containers are accessible for egg-laying and larval development (Wilke et al., 2019; Brady and Hay, 2020). In urban environments, breeding sites are usually artificial containers that accumulate water such as vessels, flower pots, discarded plastic, metallic objects, and tires (Zahouli et al., 2017; Wilke et al., 2019). In most cases, containers are filled with rainwater, which is a poor source of nutrients.

Several biotic and abiotic elements present in water are known to drive the selection of oviposition sites by gravid females. These include the presence of conspecifics and/or predators, organic matter, surrounding vegetation, moisture, salinity, ammonium, and phosphate (Wong et al., 2011; Onchuru et al., 2016; Kroth et al., 2019; Hery et al., 2021). Furthermore, microbial communities present in water-holding containers have been shown to influence *Ae. aegypti* oviposition (Benzon and Apperson, 1988; Ponnusamy et al., 2008; Melo et al., 2020). Indeed, females can locate suitable breeding sites using infochemicals emitted by microbes inhabiting the aquatic niche (Mwingira et al., 2020). This choice is probably under selection pressure because microorganisms within oviposition containers not only represent food to the larvae but can also establish intricate host-bacterial community networks eventually defining a symbiotic relation (Ponnusamy et al., 2008).

Despite the increasing interest in mosquito-microbiota interactions, the origin of the microbial communities that colonize mosquitoes and the relative contribution of the environment to their acquisition is still debated (Guégan et al., 2018; Saab et al., 2020). Several authors have shown that part of mosquito-associated bacteria is acquired during early life stages in larval habitats (Coon et al., 2014, 2016; Dada et al., 2014; Dickson et al., 2017). Besides, the bacterial communities present in *Ae. aegypti* larvae are influenced by the aquatic environment where they develop (Coon et al., 2014, 2016; Hery et al., 2021). Furthermore, some members of the bacterial community can be transstadially transmitted to adults (Lindh et al., 2008; Coon et al., 2014, 2016; Dada et al., 2014; Scolari et al., 2019; Rocha et al., 2021).

It has been suggested that mosquito females can add key microbial associates during egg-laying, affecting the microbial community within the breeding site (Coon et al., 2016). This may promote the dispersal of their associated symbionts and provide a specific microbial inoculum for offspring rather than leaving their acquisition to chance (Wong et al., 2015). Some bacteria recovered from immature stages and adults have already been detected on the eggs females lay (Coon et al., 2014; Rocha et al., 2021). Moreover, mosquitoes are capable of transferring bacteria to their oviposition sites and picking up the same bacteria from the water they emerged from (Lindh et al., 2008; Rocha et al., 2021). It has been suggested that transmission of maternal microbiota to larval breeding sites could occur directly, through egg smearing or some forms of transovarial transmission; or indirectly during egg-laying when females might unintentionally inoculate microbes into oviposition sites (Favia et al., 2007).

Although the properties of the external environment strongly regulate the bacterial communities where they are found, the dissemination of microbial cells from eukaryotic hosts can also impact the composition and traits of the microbiota in the immediate environment (Sullam et al., 2012; Wong et al., 2015). The capacity to alter the bacterial community within preadult habitats would be substantial considering the intrinsic ability of *Ae. aegypti* to exploit small and temporary water containers (Scolari et al., 2019). This ability could explain, in part, how this mosquito has successfully adapted to exploit and maximize production within confined and nutrient-scarce habitats. Interestingly, organismal environment-modifying capacities exerted by both parental individuals and their offspring during ontogenesis is a tenet of niche construction theory (Odling-Smee et al., 2013; Laland et al., 2015). Within this conceptual framework, the phenomenon of developmental niche construction can occur via chemical excretion, generation of physical structures, or dependent on the physiological properties of symbionts (Engel and Moran, 2013; Schwab et al., 2017).

Considering all the aforementioned, we intend to test the hypothesis that either microorganisms or metabolites (Mosquera et al., 2021) can be transferred by gravid female mosquitoes to shape the bacterial community of the breeding site as a strategy to enhance offspring fitness. Thus, a shift in bacterial community structure within the aquatic habitat could then be interpreted as a signature profile of oviposition and developmental activity. As such, it would become a measurable layer of evidence, suggesting developmental niche construction by gravid females and their offspring. Therefore, to address our hypothesis, we investigated whether gravid females (i) act as mechanical vectors of bacteria and (ii) modulate the bacterial community in water-holding containers through oviposition. Furthermore, we tested whether bacteria acting as oviposition indicators enhance progeny fitness.

## METHODOLOGY

### Mosquito rearing

*Aedes aegypti* (F2) were obtained from a Brazilian laboratory colony (BR URCA) established from eggs collected in ovitraps in the Urca district of Rio de Janeiro city. All mosquitoes used in the experiments were maintained under insectary conditions at 28±2°C, 70±10% relative humidity, and a 12:12 LL/DD photoperiod. Larvae were reared in plastic trays containing non-chlorinated water and fed half a tablet of TetraMin fish food (Tetra) every day. Pupae were transferred from rearing trays to cardboard cages in plastic flasks, after which adults emerged. Adults were offered 10% sucrose solution *ad libitum*. Females were blood-fed 7 days post-emergence on a Hemotek Membrane Feeding System (Hemotek Ltd) using human blood. Human blood used to feed adult mosquitoes was obtained from a blood bank (Fundação Hemominas, Belo Horizonte, MG, Brazil), according to the terms of an agreement with Instituto René Rachou, Fiocruz/MG (OF.GPO/CCO agreement-Nr 224/16). Pilot experiments revealed that this mosquito population has its oviposition peak 72 hours after a blood meal. Only fully engorged females were collected for further assays.

### Mechanical transmission of bacteria

#### Experimental design

To assess whether *Ae. aegypti* females can mechanically transfer viable bacteria to solid culture media, a single female was released in a cardboard cage (brand new, cleaned with 70% ethanol soaked paper wipes, and exposed to 15 min of UV light in a biosafety cabinet) presenting a Petri dish at the bottom loaded with either LB (Lysogeny Broth) or blood agar media. Five replicates were performed *per* culture medium tested, plus two environmental control plates *per* medium type.

After 24 hours, females were removed from the cages, pooled, and washed with 1ml of sterile phosphate-buffered saline (PBS) for 10 minutes. Moreover, a swab of the wall and bottom of the cardboard cage was collected and placed in PBS (1 ml) for 10 minutes. Subsequently, an aliquot (50 µl) of these PBS washes, both from the body surfaces and the cage swab, was inoculated on LB and blood agar plates, separately. The plates were incubated for up to 48 hours. Negative control plates with only sterile PBS resulted in no colonies.

#### DNA extraction and PCR amplification

Bacterial isolates were examined and characterized according to their features. Colonies with visually distinct morphologies were isolated from each medium, followed by total genomic DNA extraction using the DNeasy Blood & Tissue Kit (Qiagen), according to the manufacturer’s manual. A reagent blank extraction was performed as a negative control of the process.

The full-length 16S rRNA gene (∼1500pb) was amplified by the pair of primers 27F (5’-AGAGTTTGATCMTGGCTCAG-3’) and 1492R (5’-TACGGYTACCTTGTTACGACTT-3’). Polymerase chain reactions (PCR) were carried out in a 25 µL final volume using 0.50 µl of 5U/µl GoTaq Polymerase (Promega), 1.50 µl of 25 mM MgCl2, 0.50 µl of 10 mM dNTP mixture, 5 µl of 5X reaction buffer, 10 µM of each primer and 2.5 µl of template DNA. Amplification consisted of an initial denaturation at 95°C for 2 min, 30 cycles of 95°C for 30 s, 55°C for 30 s, and 72°C for 1 min 40 s, followed by a final extension at 72°C for 5 min. A PCR amplification control was performed. Reactions and negative controls were analyzed by electrophoresis in a 1% agarose gel. Controls, both from DNA extraction and PCR, showed no amplify bands.

#### Sanger sequencing and taxonomic identification

PCR products were purified using the ReliaPrep DNA Clean-up and Concentration System (Promega) following the manufacturer’s protocol. Sequencing reactions were conducted using the BigDye Terminator v3.1 Cycle Sequencing Kit (Thermo Fisher Scientific). Three primers (two forward and one reverse) were used to generate amplicons for Sanger sequencing [27F, 515F (5’-GTGCCAGCMGCCGCGGTAA-3’) and 1492R]. The combination of sequence data obtained with these three amplicons generates a contiguous sequence that encompasses most of the full 16S rRNA gene (PennVet Center for Host-Microbial Interactions, 2019). Sequencing was performed on an ABI 3730 DNA sequencer.

The sequenced reads were assembled using the software Geneious Prime (v2019.0.4). Bacterial taxonomic classification was performed using the SILVA Alignment, Classification and Tree (ACT) Service with a minimal identity with query sequences of 95%. A chord diagram representing the presence and absence patterns of the isolated bacteria from each of the groups, *per* culture medium, was plotted using the *chorddiag* package v0.1.3 in R environment.

### Changes in the bacterial profile of breeding sites

#### Experimental design

To test whether *Ae. aegypti* females modify breeding site community composition through oviposition, an experiment with five different treatments was designed. Ten replicates *per* treatment were carried out using cardboard cages presenting a plastic cup with 80 ml of type I water and 500 µl of sterilized food. This diet was prepared by dissolving finely groundfish food in type I water and autoclaving it for 20 min at 120°C. All water containers were set up on day one with sterilized food added on day 2.

Treatment 1 (T_1_) acted as an environmental control (type I water plus sterilized food). Treatment 2 (T_2_) was developed using sterilized mosquito eggs that were manually deposited. Eggs were sterilized using 70% ethanol for 5 min, followed by a wash in a 3% bleach and 0.1% benzalkonium chloride (Quatermon 30, Chemitec, Brazil) solution for 3 min, an additional wash in 70% ethanol for 5 min, and rinsing three times in sterile water. The sterile condition of eggs was confirmed by negative PCR amplification of the 16S rRNA gene V_4_ hypervariable region using the primers 515F and 806R (5’-GGACTACHVGGGTWTCTAAT-3’). Besides, this was reinforced by the absence of bacterial growth from sterilized eggs transferred to LB broth, which indicated no viable bacteria were present. Treatment 3 (T_3_) was developed using manually deposited non-sterilized eggs. Eggs, both for T_2_ and T_3_, were derived from groups of gravid females that oviposited on pieces of filter paper, which were stored under insectary conditions until needed (but not longer than one month). Treatment 4 (T_4_) was developed with a sugar-fed female that was held for 24 h without access to drinking to assure that it would interact with the water in the container and thus ensure physical contact control for mosquitoes that cannot lay eggs. Treatment 5 (T_5_) was developed with a gravid female (72 hours post-blood-feeding) that was allowed to lay eggs. For both, T_4_ and T_5_, females were removed from cardboard cages after a 24 h exposure interval. Once females were removed, the number of eggs laid in each T_5_ replicate was counted using a magnifying glass. This allowed us to calculate an average number subsequently used for manually depositing eggs in T_2_ and T_3_ replicates.

For T_2_, T_3_, and T_5_, water samples were collected when at least one pupa was detected. For T_4_ and T_1_, water samples were collected on days 15 and 16, respectively. As pupation represents a developmental checkpoint in the holometabolous cycle (Romoli et al., 2021), we considered this criterion as the basis for the sampling as it represents an environment that has successfully sustained the development of larvae. For the control (T_1_) and the treatment representing the physical interaction of an adult female mosquito and water (T_4_), we allowed the ecological succession to extend to the last days of the experiment to allow for a comparable scenario between the treatments and practical reasons when considering the DNA extraction.

#### DNA extraction and high-throughput sequencing

Each water sample was aseptically filtered through a polyethersulfone membrane (0.22 μm pore size, 50 mm diameter) using vacuum-driven filters (Biofil) and a vacuum-pressure pump (Millipore). Filter membranes were cut into small pieces using a stainless steel scalpel and placed in sterile tubes. A control using ultrapure water was carried out to verify whether the membrane or filtration process could introduce any contamination. Bacterial genomic DNA was extracted from bacterial cells retained on each filter membrane using the DNeasy PowerSoil Kit (Qiagen), following the manufacturer’s methods. A reagent blank extraction was the control of the DNA extraction process. DNA sample concentration was measured using a Qubit fluorescence assay (Invitrogen). All DNA samples were concentrated in a vacufuge concentrator (Eppendorf) and sent for amplicon sequencing (16S rRNA, V_4_ region primers) on an Illumina HiSeq PE250 instrument at Novogene Bioinformatics Technology Co., Ltd. (Beijing, China). Since the controls, both from the filtration process and from the DNA extraction, resulted in negative PCR amplification, they were not further processed and were not sequenced.

#### Bioinformatics analysis and taxonomic assignment

Raw sequence data generated were processed using the DADA2 pipeline (v1.6.0) to identify Amplicon Sequence Variants (ASVs). The raw reads were trimmed to remove the primers. The forward reads were trimmed at position 180, and reverse reads at nucleotide 150. After trimming, the reads with a maximum of 2 expected errors for the error model prediction and merging were conserved.

Taxonomic classification was assigned by TAGME (Valente Pires et al., 2018) using the pre-built model for the amplified region. HTSFilter package was used to remove ASVs containing reads less than a cutoff value defined by calculating a Jaccard index. All the above-mentioned bioinformatics tools, plus diversity and statistical analyses were executed in Rstudio v1.1.423.

#### Diversity and statistical analyses

ASVs diversity within and between samples was compared. The Simpson index (1-D) was used to measure alpha diversity. Alpha diversity metrics between groups were compared using the Kruskal-Wallis test, followed by post-hoc Dunn’s multiple comparisons tests. P-values were adjusted using the Benjamini-Hochberg method.

A Jensen-Shannon distance matrix was used for beta diversity analysis. Principal coordinates analysis (PCoA) was conducted to visualize and interpret the overall dissimilarity in the microbial community structure among the treatments. Besides that, a permutational multivariate analysis of variance (PERMANOVA) was performed to explore the significance of the presence of eggs and/or the female interaction with water, on the bacterial signatures associated with each group. A pairwise PERMANOVA was performed based on the ASVs abundance matrix transformed using the Hellinger method.

The differentially abundant ASVs were detected using DESeq2. In a multivariate model, the likelihood ratio test was used to identify differentially abundant variants. For univariate analysis, the variants differing in each variable – female interaction with water and eggs presence – were identified using the Wald test. All ASVs with adjusted P-value < 0.01 were considered differentially abundant and were used for model construction.

A general Random Forest (RF) model was built using all the previously identified ASVs and the Gini importance of each ASV was calculated. The 30 most important ASVs were used to construct models for each variable – female interaction with water and egg presence. 1000 bootstrap analyses were performed by randomly selecting 50% of samples from the analyzed variable, building 100 trees, and calculating the importance of each ASV. The 10 most important ASVs among the 1000 tests were chosen to build a final predictive model. The model construction and performance analysis were executed in R environment using *caret* package v6.0-86.

As the predictive model tested by the RF approach identified features capable of discriminating the communities based on the key experimental variables, we deemed it relevant to search for indicator taxa. This ecological analysis was executed to identify ASVs that reflect the effects that biotic and/or abiotic factors, encompassed within each treatment, exert, thus shaping the community composition. In particular, we aimed to identify ASVs whose occurrence and abundance provide evidence of the impact that oviposition and larval development (T_5_) had upon the breeding site bacterial consortium. The analysis was performed in R environment using the *indicspecies* package v1.7.7.

Finally, we used a modified Raup–Crick metric (Bray-Curtis-based Raup-Crick, RCbray) to assess whether the compositional turnover detected, particularly between T_1_ (control) and T_5_ (female oviposition), was driven primarily by drift as the force structuring the assembled communities. The beta net relatedness index (betaNRI) was used with the RCbray to determine the presence of a pattern due to ‘basal’ phylogenetic turnover of lineages caused by replacement. All statistical analyses were performed in R environment using the *microeco* package v0.3.3.

### Effects of *Elizabethkingia* on larval development, mortality, and adult size

#### Selection of bacteria for larvae fitness experiments

To assess if ASVs identified as an indicator of oviposition activity may have an impact on *Ae. aegypti* development, a bacterial strain belonging to the genus *Elizabethkingia* was selected for fitness experiments. As a control, we also tested *Asaia*, a bacterium widely present in the microbiota of several mosquito species.

*Asaia* sp. strain AE06 (GenBank accession: KR703670) was recovered from the midgut of adult females of the Paea laboratory strain, which was established in 1994 (Vazeille-Falcoz et al., 1999). *Elizabethkingia* sp. strain VV01(GenBank accession: KU096882) was isolated from field-collected mosquitoes. Wild *Ae. aegypti* were collected in Colônia Z-10 (22°49’23.50”S; 43°10’42.93”W), a fishermen community in Rio de Janeiro. Larvae, water, and deposited sediment were collected from two natural breeding sites, brought to the insectary, and conditioned in clean disposable cups at 27±2°C. No additional food or water was added to the cups until adult emergence. Adults were fed *ad libitum* with sterilized cotton soaked in sterilized 10% sucrose solution until midgut dissection. Ice-anesthetized adult female mosquitoes were surface-sterilized in 70% ethanol for 1 min and rinsed in sterile PBS. As surface sterilization control, individuals were rinsed in sterile PBS, which was plated on LB plates. Midguts were removed over a sterile glass slide and macerated in sterile PBS. Each midgut sample was 10-fold diluted and plated on LB and tryptone soy agar plates. For the next 72h, bacteria were screened based on colony morphology. Samples from each different bacterial morphotype were preserved and stored at -70ºC. Bacterial DNA was extracted by a conventional boiling and freezing step. A 16S rRNA gene segment between the V_1_-V_3_ hypervariable regions was amplified by PCR using the primers 27F (5’-AGAGTTTGATCCTGGCTCAG-3’) and 536R (5’-GTATTACCGCGGCTGCTG-3’) and Sanger sequenced for taxonomic identification.

#### Experimental design

At 24 hours post-hatching 36 L_1_ larvae (Paea strain) *per* group were individually placed in the wells of three 12-well cell culture plates. Each well received 4 ml of non-chlorinated water, 3 mg of TetraMin fish food, and 100μl of *Asaia* or *Elizabethkingia* culture suspended in PBS (OD600=1). Controls received 100μl of PBS. Larvae development was monitored three times a day (8:00, 12:00, and 17:00) to record mortality and molt for each insect. Developmental time was monitored up to the day all immatures reached the adult stage or died. The wing length was measured, excluding the fringe, as a proxy for adult body size (Harbach and Knight, 1980). During experiments, specimens were maintained at 27±2°C and 70±10% relative humidity. Larvae were not antibiotic-treated before bacteria exposure. To verify *Asaia* and *Elizabethkingia* colonization in the larval guts, L_4_ midguts were dissected, homogenized in PBS, and plated on LB plates and *Asaia* specific isolation medium (AIM) (Yamada et al., 2000). Isolated bacterial strains were taxonomically identified using the 16S rRNA gene sequencing procedure previously mentioned.

#### Statistical analysis

The non-parametric Kaplan-Meier survival analysis was performed to assess whether exposure to *Asaia* or *Elizabethkingia* affected the duration of larval instars (L_1_, L_2_, L_3_, and L_4_) and the pupal stage. The total immature development time (L_1_ to adult) and larval survival were also compared between *Ae. aegypti* groups using the same approach. Global differences between the control, *Asaia*, and *Elizabethkingia*-exposed larvae were tested by the Log-rank test. When significant variation was detected, paired tests were then performed. Significance levels were corrected by the Bonferroni criterion. Wing lengths were compared using the Kruskal-Wallis test. Statistical analyses were carried out using R (v3.2.3).

## RESULTS

### *Aedes aegypti* females transmit bacteria mechanically

Our results demonstrated that *Ae. aegypti* females transfer culturable viable bacteria to solid culture media (Figure 1). To further dissect the possible sources of these bacteria, a cage swab, mosquito body washes, and environmental controls were performed for each medium tested.

**Figure 1.**
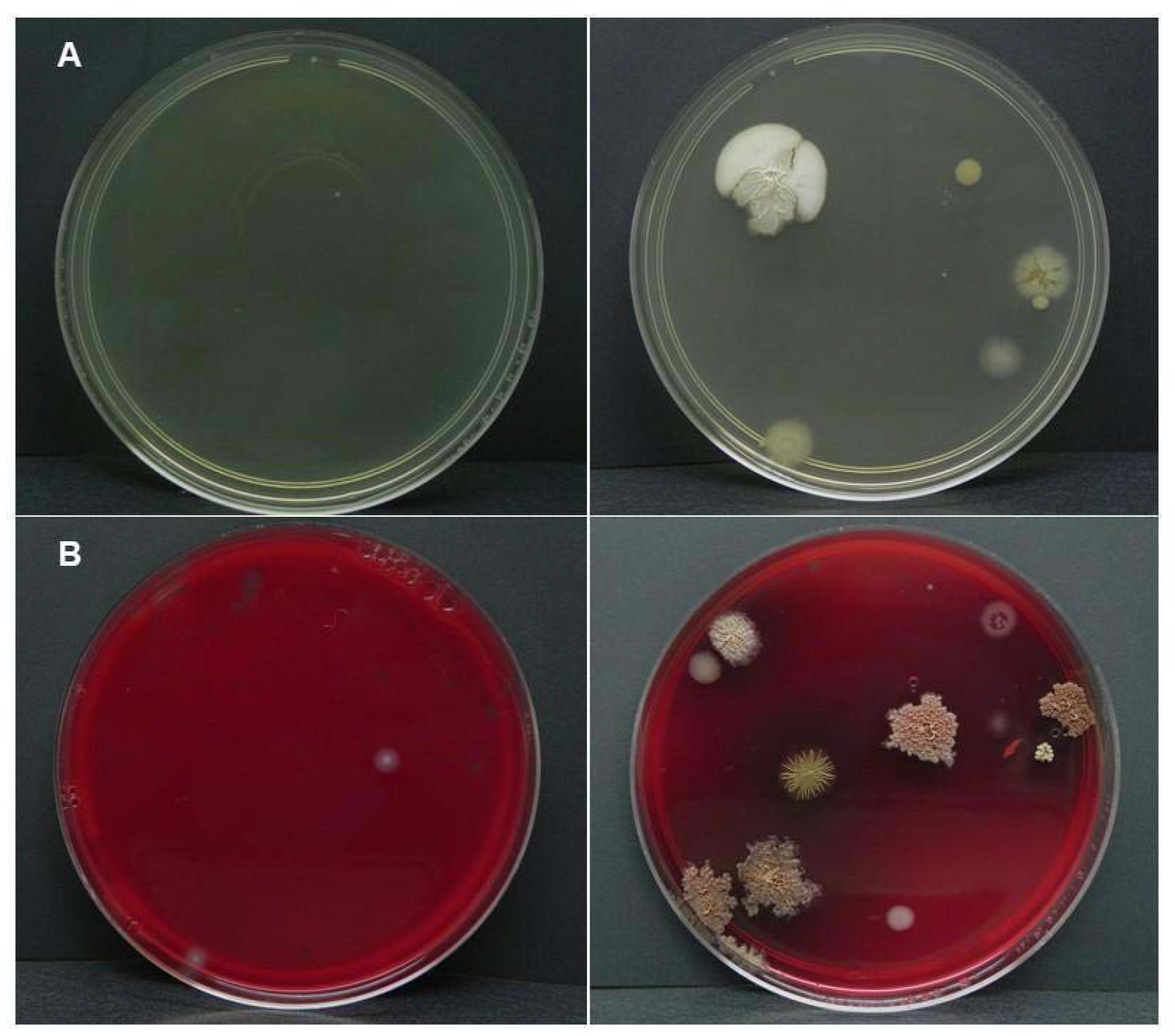
*Aedes aegypti* females transfer viable and cultivable bacteria to solid culture media. Growth of bacterial colonies on LB (A) and Blood agar (B). On the left, environmental control plates, and on the right, plates exposed to interaction with a gravid female.

A total of 28 isolates were recovered from LB plates (Figure 2). Altogether, these isolates belonged to three phyla, six families, and seven genera (Supplementary material 1). The bacterial diversity observed in LB plates exposed to interaction with a gravid female mosquito was notably higher compared to that seen in control plates (Figure 1). Bacteria isolates recovered from cage swab plates were assigned to the genus *Serratia*. Bacterial isolates from body washes from gravid female mosquitoes were classified into genera *Serratia* and *Elizabethkingia. Bacillus* was the most common genus of bacteria found after interaction with a gravid female, and together with *Ornithinibacillus, Lysinibacillus*, and *Kroppenstedtia* constituted the genera exclusively associated with this experimental condition. Besides, the genera *Serratia* and *Paenibacillus* were also reported from female-exposed LB plates. The environmental controls showed bacterial growth in one of the two plates examined. This isolate was assigned to the genus *Paenibacillus* (Figure 2).

**Figure 2.**
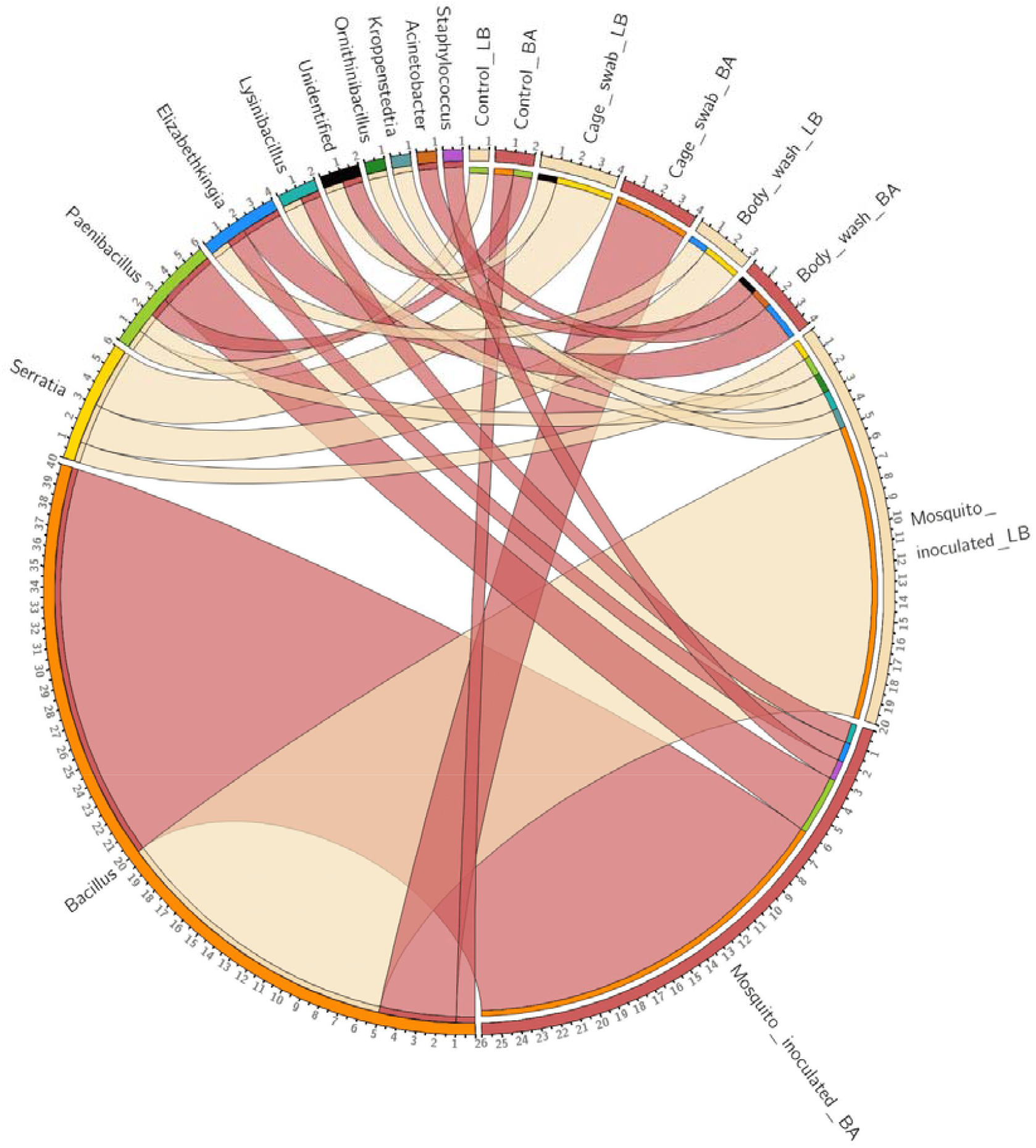
Chord diagram showing the bacterial genera recovered from LB (beige) and blood agar plates (red) and the different sources from which these were isolated.

Culturable bacteria isolated from blood agar plates were represented by 36 isolates (Figure 2). Bacteria belonged to three phyla, seven families, and six genera (Supplementary material 2). Similarly, as observed with the LB medium, bacterial isolates recovered from blood agar plates exposed to a gravid female mosquito showed higher diversity compared with those from environmental controls (Figure 1). Isolates obtained from the cage swab were members of the genus *Bacillus*. Blood agar plates on which the body wash of gravid females was plated generated four bacterial isolates assigned to genera *Elizabethkingia* and *Acinetobacter*. As with LB plates, *Bacillus* was the most common genus reported in blood agar plates visited by mosquitoes. The genera *Lysinibacillus* and *Staphylococcus* were exclusively associated with female visited samples. Besides, *Paenibacillus* and *Elizabethkingia* were also isolated in this condition. Finally, bacterial isolates recovered from the environmental controls were assigned to the genera *Bacillus* and *Paenibacillus*.

It is important to stress that *Serratia* (LB) and *Elizabethkingia* (blood agar) were the only genera shared between plates exposed to interaction with a gravid female and those from mosquito body washes.

### Changes in the bacterial profile of the breeding site

High throughput sequencing targeting the V_4_ region of the bacterial 16S rRNA gene produced 20,425,105 reads from 50 water samples. After computational quality control, 16,896,903 reads were considered for taxonomic analysis. A data matrix was generated encompassing 532 ASVs. Nonetheless, the HTSFilter package identified a cutoff of 75 reads based on the Jaccard index. Therefore, all ASVs below this value were removed for downstream analysis. The total number of ASVs identified above the cut-off value was 159.

The alpha diversity was significantly different between experimental groups (Kruskal-Wallis, P = 0.01). Furthermore, the post-hoc Dunn test identified that T_5_ had a significantly lower Simpson index (1-Simpson) compared with the other four treatments (Supplementary material 3). Water samples belonging to T_2_ had the highest ASV diversity (mean Simpson index = 0.768), while T_5_ presented the lowest one (mean Simpson index = 0.527) (Figure 3).

**Figure 3.**
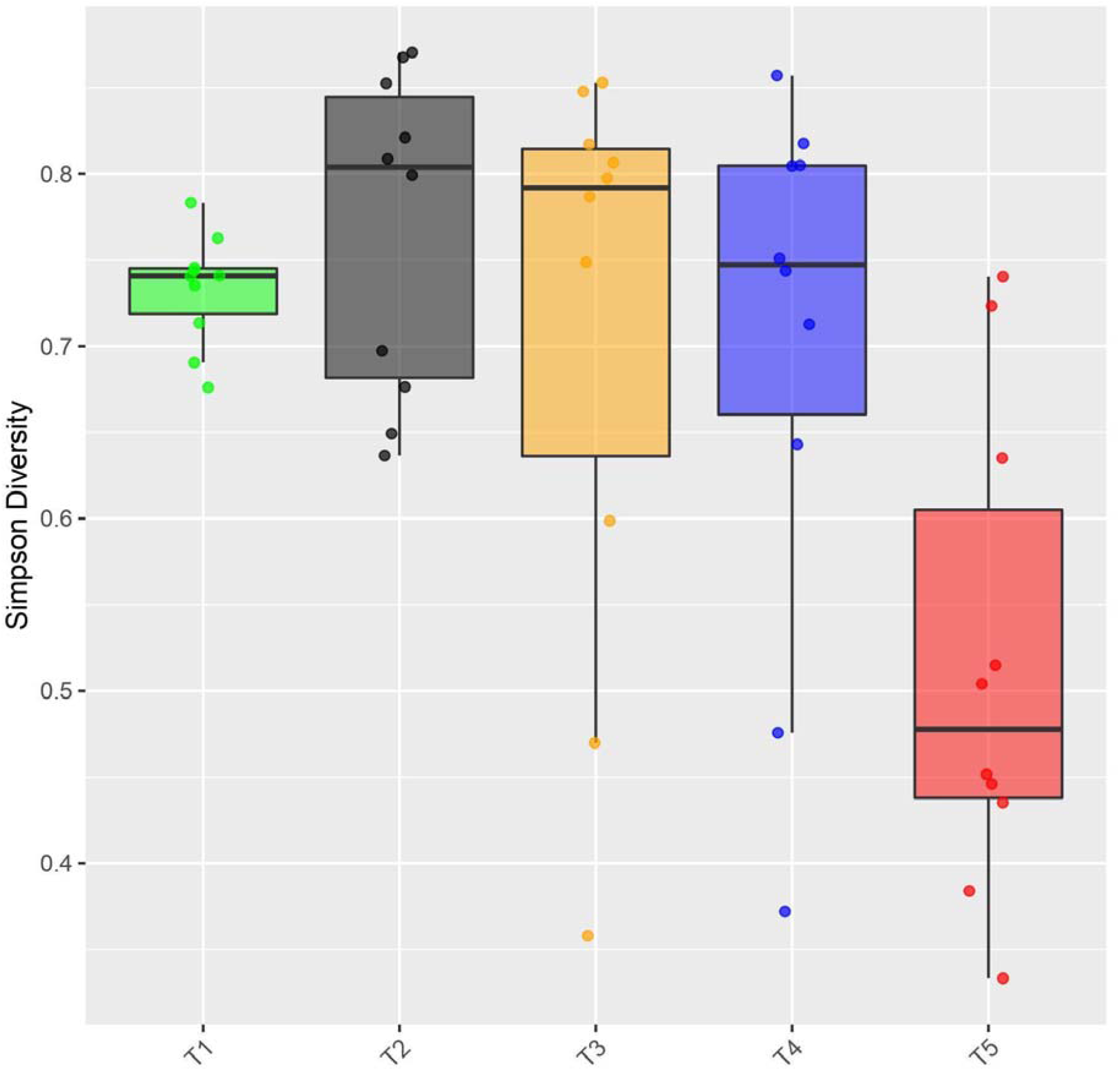
Alpha diversity of ASVs as a function of treatment. Comparison of Simpson’s Index of Diversity (1-Simpson’s Index) recorded for the different treatments using a boxplot (10 replicates and median). The alpha diversity was significantly different between experimental groups (Kruskal-Wallis, P = 0.01). Treatment 5 had a significantly lower Simpson index compared with the other four treatments.

The Jensen-Shannon divergence metric was used to compare ASV diversity among treatments (Supplementary material 4). The PCoA captured around 48% of the variation in Jensen-Shannon distance along the two chosen axes (PCo_1_ and PCo_2_) represented in Figure 4. A comparison of the bacterial communities associated with each treatment showed distinct clustering patterns. Samples belonging to T_3_ and T_5_ displayed higher inter-treatment variability clustering bottom and top right, respectively (Figure 4).

**Figure 4.**
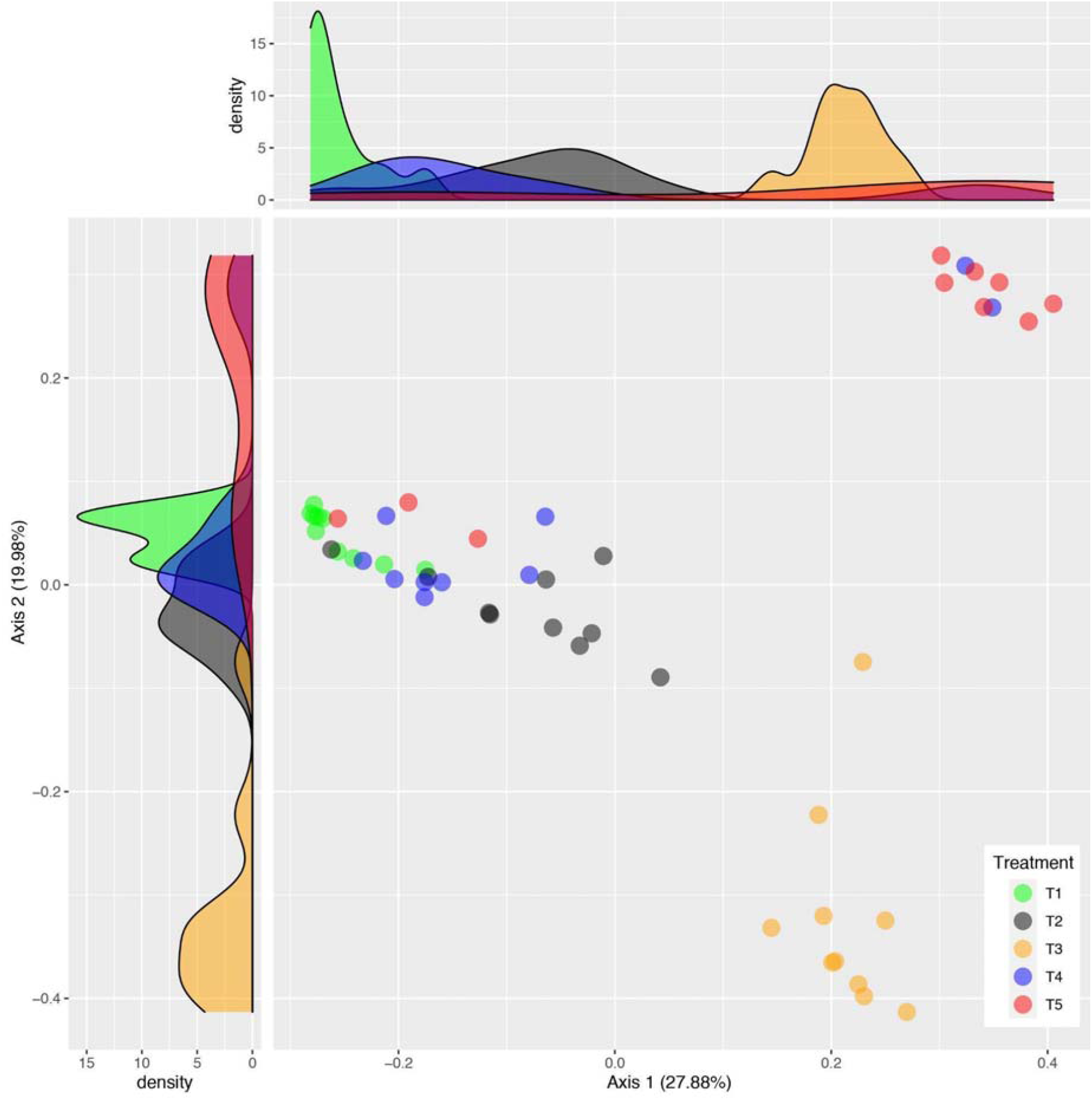
Beta diversity analysis. Principal coordinate analysis (PCoA) of Jensen-Shannon distances. Distinct clustering patterns for each experimental treatment and their corresponding replicates are represented by a color code. Axis 1 (27.88%) and axis 2 (19.9%) show the percentage of variation explained.

PERMANOVA revealed that presence of eggs (R^2^ = 0.206, df = 2, P = 0.001), female interaction with water (R^2^ = 0.103, df = 1, P = 0.001), and the interaction of these two variables (R^2^ = 0.092, df = 1, P = 0.001) explain 40% of the variance in bacterial composition over the groups of samples. Additionally, the pairwise PERMANOVA confirmed that all groups differ significantly from each other in terms of beta diversity, suggesting that there are consistent bacterial signature profiles for each condition (Supplementary material 5).

The RF model selected 10 ASVs as the most important features for each variable explored – female interaction with water and egg presence (Tables 1 and 2). The bacterial signatures modeled by RF had robust prediction performances supported by their high AUC values (Supplementary material 6).

**Table 1.** ASVs predictive of female interaction with water. The 10 most important discriminating ASVs identified by RF when a female *Aedes aegypti* interacted with water.

| ASV ID | Family | Species | MeanDecreaseGini |
| --- | --- | --- | --- |
| sq2 | Weeksellaceae | *Elizabethkingia anophelis* | 2.7596470 |
| sq81 | Xanthomonadaceae | *Stenotrophomonas maltophilia* | 1.8039565 |
| sq12 | Rhizobiaceae | *Agrobacterium radiobacter* | 1.0807458 |
| sq10 | Sphingomonadaceae | unclassified_Novosphingobium | 1.0666743 |
| sq23 | Burkholderiaceae | unclassified_Pseudacidovorax | 0.9449007 |
| sq21 | Enterobacteriaceae | unclassified_Enterobacter | 0.8477820 |
| sq6 | Moraxellaceae | unclassified_Acinetobacter | 0.8216041 |
| sq28 | Xanthomonadaceae | *Stenotrophomonas maltophilia* | 0.7735763 |
| sq49 | Burkholderiaceae | *Herbaspirillum huttiense* | 0.6960721 |
| sq9 | Pseudomonadaceae | *Pseudomonas putida* | 0.7169478 |

**Table 2.** ASVs predictive of egg presence. The 10 most important discriminating ASVs identified by RF when *Aedes aegypti* eggs were present in water.

| ASV ID | Family | Species | MeanDecreaseGini |
| --- | --- | --- | --- |
| sq31 | Burkholderiaceae | unclassified_Pelomonas | 2.3970572 |
| sq45 | Xanthobacteraceae | *Bradyrhizobium japonicum* | 2.3434994 |
| sq1 | Caulobacteraceae | *Caulobacter vibrioides* | 2.2397369 |
| sq11 | Burkholderiaceae | *Ralstonia insidiosa* | 1.6738255 |
| sq67 | Nocardiaceae | *Rhodococcus erythropolis* | 1.3978588 |
| sq18 | Sphingomonadaceae | unclassified_Sphingomonas | 1.3206670 |
| sq42 | Xanthobacteraceae | *Afipia genosp.* | 1.2906533 |
| sq23 | Burkholderiaceae | unclassified_Pseudacidovorax | 0.9296086 |
| sq2 | Weeksellaceae | *Elizabethkingia anophelis* | 0.9353752 |
| sq10 | Sphingomonadaceae | unclassified_Novosphingobium | 0.8482134 |

On the other hand, indicator species analysis identified ASVs considered to be specific microbial features associated with the act of oviposition and larval development, i.e. treatment 5. Seven ASVs were pinpointed as oviposition-indicating species as they possess significant fidelity and predictive value towards the ecological conditions represented in this niche/treatment (Table 3). Indicator species were assigned to the following taxa: *Leifsonia soli, Elizabethkingia anophelis, Paenibacillus polymyxa, Stenotrophomonas maltophilia, Elizabethkingia, Methylobacterium*, and *Elizabethkingia meningoseptica*.

**Table 3.** Indicator species analysis. ASVs considered features of oviposition activity (treatment 5).

| ASV ID | Bacterial taxa | Specificity | Fidelity | Indicstat | P-value |
| --- | --- | --- | --- | --- | --- |
| sq44 | *Leifsonia soli* | 0.8540 | 0.9000 | 0.877 | 0.001 |
| sq2 | *Elizabethkingia anophelis* | 0.7274 | 1.0000 | 0.853 | 0.001 |
| sq75 | *Paenibacillus polymyxa* | 0.7911 | 0.8000 | 0.796 | 0.001 |
| sq81 | *Stenotrophomonas maltophilia* | 0.5068 | 0.9000 | 0.675 | 0.004 |
| sq86 | *Elizabethkingia* | 0.7355 | 0.6000 | 0.664 | 0.001 |
| sq85 | *Methylobacterium* | 0.7138 | 0.6000 | 0.654 | 0.034 |
| sq143 | *Elizabethkingia meningoseptica* | 0.9362 | 0.4000 | 0.612 | 0.003 |

The null model analysis employing RCbray and βNRI dissimilarity matrices showed that the main processes driving the assembly of the compared communities were: drift > homogeneous selection > dispersal limitation (Table 4).

**Table 4.** The percentage assigned to each of the processes driving the differences in community structure between the control treatment and the oviposition treatment.

| Community ecology processes | Percentage |
| --- | --- |
| Variable selection | 0.00 |
| Homogeneous selection | 13.69 |
| Dispersal limitation | 0.52 |
| Homogeneous dispersal | 0.00 |
| Drift | 85.79 |

### *Aedes aegypti* exhibits faster development in the presence of *Elizabethkingia*

Mean total immature development (L_1_ to adult) took 180.87, 173.23, and 171.04 hours for the control, *Asaia*, and *Elizabethkingia*-exposed larvae, respectively. There were no significant differences in the duration of immature development between *Asaia*-exposed *Ae. aegypti* and those reared under control conditions (Figure 5, Supplementary material 7 and 8). On the other hand, *Elizabethkingia*-exposed larvae developed faster from L1 to L2 (Chisq = 11.8, P-value < 0.01) and also from L1 to the adult stage (Chisq = 12.5, P-value < 0.01). Regarding survival, control, *Asaia* and *Elizabethkingia* exposed specimens presented 15, 8, and 23% of mortality, respectively. Survival (Figure 5, Supplementary material 7) and wing length (KW chi-squared = 0.3, P > 0.05) were not considered different among the three larval groups.

**Figure 5.**
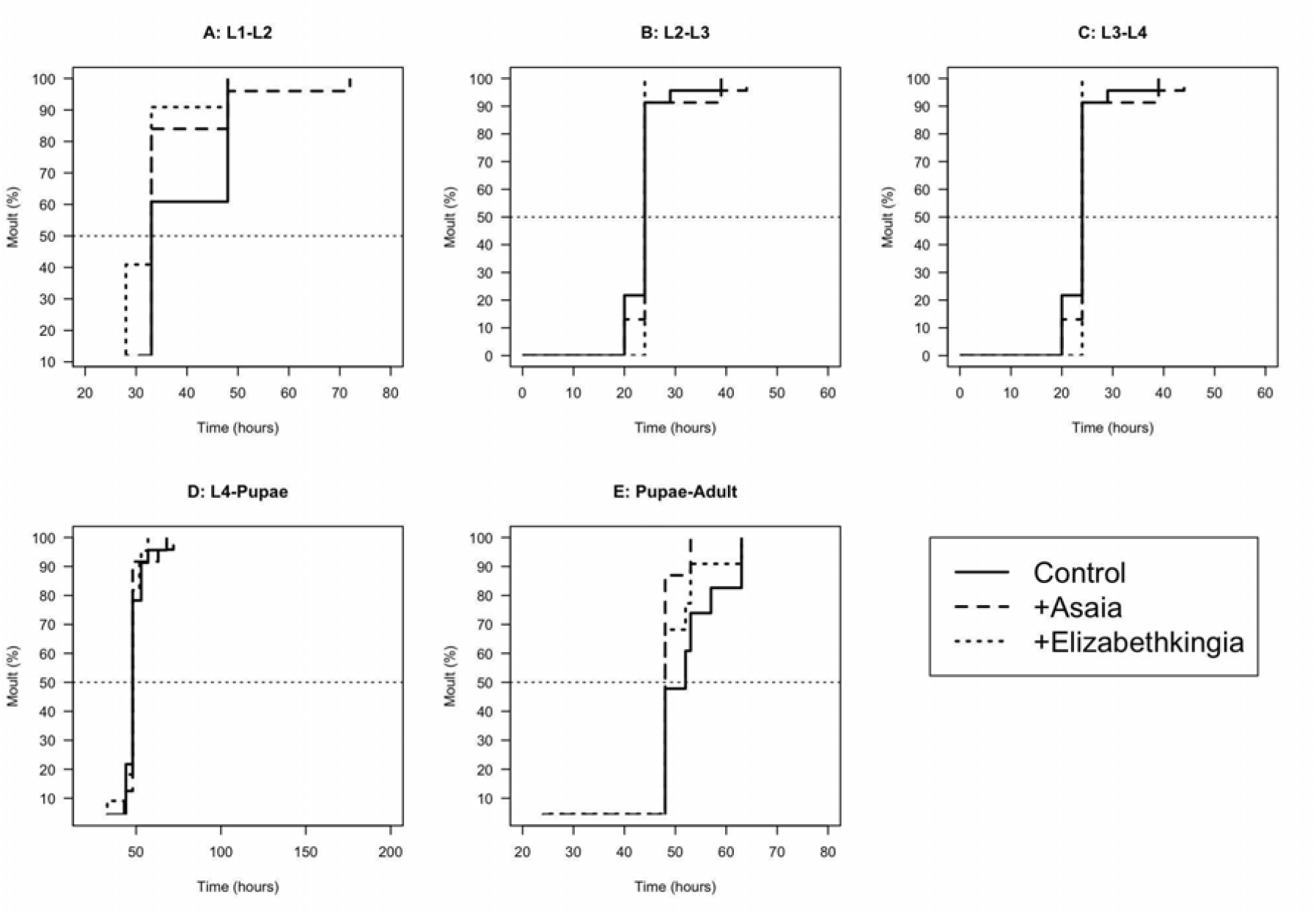
Duration of larval stages (graphs A to D) and pupal phase (E) of immature forms *Aedes aegypti* exposed to bacteria of the genera *Asaia* (+ Asia) and *Elizabethkingia* (+ *Elizabethkingia*) and the control group.

The diversity of culturable microbiota was variable in all three groups, with only the *Bacillus* genus being ubiquitous to all conditions (Supplementary material 9). *Asaia* was not isolated from any of them, while *Elizabethkingia* was recovered from the midgut of larvae exposed to it.

## DISCUSSION

In this study, we have explored the hypothesis that ovipositing females could influence the microbial consortium of the aquatic niche, which could eventually shape the microbial community within the breeding site (Coon et al., 2016), promoting larval fitness. Our initial results showed that gravid females were capable of transmitting viable and culturable bacteria already reported as mosquito symbionts. We then demonstrated that the act of oviposition promoted a significant decrease in the bacterial diversity found in breeding sites. Furthermore, this was also associated with a specific bacterial profile which included a series of indicator taxa linked to female oviposition. We present evidence that demonstrates that one of these taxa, i.e. *Elizabethkingia*, was able to accelerate larval development. Altogether, these results seem indicative of niche construction in *Aedes aegypti* breeding sites (Schwab et al., 2017).

Experiments showing mechanical transmission from females to plates confirm that gravid mosquitoes can inoculate diverse bacteria to culture media. The most predominant bacteria identified from plates visited by a gravid female (for both blood agar and LB media) was *Bacillus*. This bacterial taxon has been already identified in stable association with larvae (Luxananil et al., 2001) and the crop of adult *Ae. aegypti* (Gusmão et al., 2007). Interestingly, several bacteria inoculated belonged to strains frequently reported as key members of mosquito microbiota, e.g. *Elizabethkingia* and *Serratia* (Steven et al., 2021). Indeed, both bacterial taxa, vertically, horizontally, and transstadially transmitted (Lindh et al., 2008; Chen et al., 2015; Rocha et al., 2021), have been implicated in female sugar digestion processes (Gusmão et al., 2007; David et al., 2016), and seem fit to survive the redox conditions generated by the catabolism of blood (Muturi et al., 2018). It should be noted that moist agar plates eventually induced oviposition. Therefore, we suggest that while exploring a tentative oviposition site, gravid females inoculate the substrate with bacterial partners that according to our experiments support offspring development.

Next, we evaluated whether gravid females influence the bacterial communities of water-holding containers during egg-laying. Our findings showed that the act of oviposition, represented by treatment 5, significantly decreased the diversity of the bacterial communities present in the breeding sites. As the Simpson index is a dominance metric (Kim et al., 2017), an uneven microcosm (dominated by the most abundant taxa) could represent an advantageous scenario for larvae, as not all microbes are beneficial, whether because they are pathogenic or because they are not fulfilling key functions in the host-microbe network (Foster et al., 2017). As a substantial proportion of the microbiota associated with immature stages is acquired from the aquatic niche, an ecosystem with a low-diversity microbial community but high-fidelity microbial partners would favor the establishment of specific mutualistic interactions (Reese and Dunn, 2018). The above is congruent with recent observations made by Martinson and Strand (2021), who highlighted the successful development of *Ae. aegypti* larvae in low diversity (gnotobiotic communities) breeding sites given certain dietary conditions were met. Additionally, a decrease in evenness within a community, as seen in T_5_, can be considered a hallmark sign of susceptibility towards the establishment of an invading organism, in this case, the larvae, in the community (Wittebolle et al., 2009; Daly et al., 2018).

Regarding beta diversity, we observed significant differences in the microbial signature profiles associated with each experimental group. The ordination analysis (Figure 4) showed how the community structures diverge among treatments, particularly highlighting differences driven by uncoupling mechanical aspects of oviposition. As each treatment represents potential sources of microbial inocula, it is relevant to highlight how the single unit of the natural egg-laying and larval development process (T_5_) presented a predominant profile diverging from the others in the ordination space. Similarly, non-sterile eggs, representing the general mechanism by which mosquito colonies are reared in insectary conditions, produced a significantly different profile at the endpoint measured as the developmental checkpoint of pupation. Whether these differing bacterial environments in which the larvae of both treatments developed influence adult life-history traits could be taken into account when elaborating experimental designs evaluating the role of the microbiome under controlled conditions. It is also relevant to observe that surface-sterilized eggs manually deposited in the water presented a profile resembling that of the control and non-gravid female-water interactions. Thus we propose that stereotypical female behaviors expressed while ovipositing (e.g. grooming, tasting, defecating) would be fundamental in producing the bacterial profile represented by treatment 5, more so given that larval presence on its own (foraging in the water column and their physiological outputs) did not lead to similar profiles of beta diversity under equal conditions.

Since we identified several taxonomic indicator units belonging to the genus *Elizabethkingia* (sq2, sq86, and sq143) and that *Elizabethkingia* (sq2) also appeared as a significant bacterial feature according to the RF model, we suggest that it plays a major role in breeding site development. *Elizabethkingia* spp. has attracted much interest due to its close association with mosquitoes (Steven et al., 2021), and several of its genes seem to relate to sugar transport and metabolism, erythrocyte lysis, and protection against bloodmeal-induced oxidative stress (Kukutla et al., 2014). Besides, mosquitoes colonized experimentally with *Elizabethkingia* in their gut produce more eggs, i.e., the bacteria positively influence the fecundity of the host (Chen et al., 2020). All these observations strongly suggest a symbiotic relationship between *Elizabethkingia* and mosquitoes.

*Elizabethkingia* was shown to be capable of inhibiting *Pseudomonas*, another mosquito-associated bacterium, via an antimicrobial independent mechanism (Ganley et al., 2020). Indeed, *Elizabethkingia* has broad antibiotic resistance because of a large number of genes encoding efflux pumps and β-lactamases present in its genome (Kukutla et al., 2014). Bacteria use diverse mechanisms to compete with other members of the microbial community (Foster et al., 2017), and based on the above information, it is very likely that *Elizabethkingia* disturbs the bacterial consortium by eliminating competitors and determining which strains persist at the breeding site, as our experiment seems to suggest.

To better understand the compositional turnover reported here, the null model analysis (Stegen et al., 2013; Liu et al., 2017) revealed that at the endpoint (time of measuring the beta diversity between treatments 1 and 5) the community structures detected were not only the result of a stochastic process (drift) but also the outcome of the deterministic ecological features acting as shaping forces of diversity (Table 4). A more in-depth study of how this occurs, while considering the theoretical implications of selection and dispersal forces, is enticing and could be pursued to better understand the dynamics involved in the ecological succession within the environment where mosquitoes develop. For instance, when does the diversity shift occur? Which mechanisms are involved in rendering the community non-resistant to the disturbance, or unable to be resilient and reacquire its original structure through time? All conceptually relevant questions for disturbance ecology applied to microbiomes (Christian et al., 2015).

Finally, we evaluated whether *Elizabethkingia* and *Asaia* influenced larval developmental time, survival, and adult size in *Ae. aegypti*. This was intended to compare the impact of this oviposition-indicating taxon with that of *Asaia*, another bacterium reported in mosquito microbiota but not found in our breeding sites. Contrary to what was observed for *Anopheles gambiae*, for which *Asaia* accelerates larval development (Mitraka et al., 2013), this bacterium did not show significant effects on *Ae. aegypti*. Conversely, *Elizabethkingia* significantly speeded up development and colonized the larval midgut, which was not observed for *Asaia*, suggesting a facilitated interaction between them. Indeed, reducing mosquito larval development time might increase the probability of reaching adulthood (Díaz-Nieto et al., 2016). In this context, the presence of *Elizabethkingia* in breeding water and larval midguts has likely aided metabolic activities, providing nutrients or metabolites that stimulate faster larval development and/or represented an additional source of food (Coon et al., 2014; Chen et al., 2015). Taken together our results suggest that females can spike breeding sites with this symbiotic bacterium to support offspring fitness, which could be interpreted as a form of niche construction.

Therefore, by using a set of different experimental and analytical methodologies, and testing field-originated bacterial symbionts, we revealed an ecological and functional connection between egg-laying activity, bacterial communities in *Ae. aegypti* breeding sites, a key symbiotic bacterial taxon, and the speed of larval development. There are other microorganisms with individual or combined positive impacts on the larval development of *Ae. aegypti* (Valzania et al., 2018; Martinson and Strand, 2021). As such, we concur with the concept that these effects most likely will be understood from a community ecology perspective (Steven et al. 2021). Other indicator species from our set, as well as other microorganisms and their interactions, could be the driving forces detected in the compositional profile of T_5_ and be key to the success of the breeding site.

Niche construction theory recognizes that organisms can modify both biotic and abiotic components of their environments. This process is an outcome of their activities, metabolism, and choices, and its main consequence is to increase survival probabilities (Laland and O’Brien, 2011; Odling-Smee et al., 2013). Altogether, our findings provide evidence that an indicator taxa linked to the full act of oviposition, revealed by the modified community structure of the breeding site water, positively affects offspring fitness, strengthening the possibility of niche construction as a strategy of female *Ae. aegypti* to disseminate symbiotic bacteria through egg-laying to grant proper environments for their progeny. As stressed by Schwab and collaborators (2017), our results are in agreement with niche construction theory criteria: a substantial environment modification was detected (bacterial community diversity), and positive fitness/developmental consequences were measured when a biomarker taxon was used as an effector. These findings provide solid grounds to build upon and improve our knowledge regarding how endo and ecto microbiomes may be critical when addressing their links to host phenotypes through the lens of niche construction theory. Other layers of information may be relevant to improve this take, as recently reported by Mosquera and collaborators (2021), the metabolite profile of breeding sites also reflects the act of oviposition and development. Disentangling whether and how individual microorganisms, or their networks, exert effects on mosquito life-history traits is a growing field of study benefiting from the synergy of microbiology, ecology, and computational biology.

## DATA AVAILABILITY

Raw sequence data are available at the European Nucleotide Archive (ENA), project number: PRJEB51063.

## AUTHOR CONTRIBUTIONS

All authors contributed to the study conception and design. KMD wrote the initial draft. KMD, LEMV, MRD, RMF, and MGL critically reviewed the manuscript. KDM, LEMV, and MRD performed the experiments. KDM, LEMV, GRF, and MRD conducted data analysis. RMF, LAM, and MGL acquired funding. All authors read and approved the final manuscript.

## ACKNOWLEDGMENTS

We would like to thank the René Rachou Institute Sequencing Platform for kindly allowing us to submit our samples for Sanger sequencing. We would also like to thank Assmaa El Khal and Bianca Daoud for their technical assistance in molecular biology procedures and mosquito rearing, respectively.

