## Supplementary Materials for "Egg-laying by female *Aedes aegypti* shapes the bacterial communities of breeding sites"

**SUPPLEMENTARY MATERIAL**

**Supplementary material 1.** Bacterial taxonomic affiliation of isolates recovered from LB agar plates.

| **Plate where the isolate came from** | **Isolate** | **No. of bases used to establish identity** | **Phylum** | **Presumable genus (% sequence identity)** |
| --- | --- | --- | --- | --- |
| Cage swab | L1 | 916 | Proteobacteria | *Serratia* (100) |
| Cage swab | L2 | --- | --- | unidentified |
| Cage swab | L3 | 885 | Proteobacteria | *Serratia* (99.7) |
| Cage swab | L4 | 914 | Proteobacteria | *Serratia* (99.6) |
| Body wash 1 | L5 | 916 | Proteobacteria | *Serratia* (99.7) |
| Body wash 2 | L6 | 709 | Bacteroidetes | *Elizabethkingia* (95.4) |
| Body wash 2 | L7 | 783 | Proteobacteria | *Serratia* (94.9) |
| Female 1 | L8 | 1339 | Firmicutes | *Bacillus* (95.8) |
| Female 1 | L9 | 828 | Firmicutes | *Bacillus* (99.5) |
| Female 1 | L10 | 840 | Firmicutes | *Bacillus* (99.5) |
| Female 1 | L11 | 836 | Firmicutes | *Ornithinibacillus* (98.3) |
| Female 2 | L12 | 1328 | Firmicutes | *Bacillus* (98.8) |
| Female 2 | L13 | 1421 | Firmicutes | *Bacillus* (99.9) |
| Female 3 | L14 | 846 | Proteobacteria | *Serratia* (99.7) |
| Female 3 | L15 | 910 | Firmicutes | *Paenibacillus* (98.7) |
| Female 4 | L16 | 783 | Firmicutes | *Bacillus* (99.6) |
| Female 4 | L17 | 1372 | Firmicutes | *Bacillus* (98.5) |
| Female 4 | L18 | 1333 | Firmicutes | *Lysinibacillus* (97.6) |
| Female 4 | L19 | 1330 | Firmicutes | *Bacillus* (98.5) |
| Female 4 | L20 | 1340 | Firmicutes | *Bacillus* (98.2) |
| Female 4 | L21 | 829 | Firmicutes | *Bacillus* (97.9) |
| Female 4 | L22 | 1376 | Firmicutes | *Kroppenstedtia* (92.4) |
| Female 5 | L23 | 832 | Firmicutes | *Bacillus* (99.2) |
| Female 5 | L24 | 833 | Firmicutes | *Bacillus* (99.6) |
| Female 5 | L25 | 823 | Firmicutes | *Bacillus* (99.7) |
| Female 5 | L26 | 1367 | Firmicutes | *Bacillus* (99.9) |
| Female 5 | L27 | 1373 | Firmicutes | *Bacillus* (98.4) |
| Control 1 | L28 | 1373 | Firmicutes | *Paenibacillus* (99.8) |

**Supplementary material 2.** Bacterial taxonomic affiliation of isolates recovered from blood agar plates.

| **Plate where the isolate came from** | **Isolate** | **No. of bases used to establish identity** | **Phylum** | **Presumable genus (% sequence identity)** |
| --- | --- | --- | --- | --- |
| Cage swab | B1 | 1361 | Firmicutes | *Bacillus* (100) |
| Cage swab | B2 | 1332 | Firmicutes | *Bacillus* (100) |
| Cage swab | B3 | 1329 | Firmicutes | *Bacillus* (100) |
| Cage swab | B4 | 1298 | Firmicutes | *Bacillus* (99.7) |
| Body wash 1 | B5 | 1412 | Bacteroidetes | *Elizabethkingia* (99.7) |
| Body wash 1 | B6 | 1338 | Proteobacteria | *Acinetobacter* (98) |
| Body wash 2 | B7 | 794 | Bacteroidetes | *Elizabethkingia* (99.8) |
| Body wash 2 | B8 | --- | --- | unidentified |
| Female 1 | B9 | 1355 | Firmicutes | *Paenibacillus* (100) |
| Female 1 | B10 | 1390 | Firmicutes | *Bacillus* (99.9) |
| Female 1 | B11 | 1376 | Firmicutes | *Bacillus* (99.9) |
| Female 1 | B12 | 1348 | Firmicutes | *Bacillus* (99.6) |
| Female 2 | B13 | 1423 | Firmicutes | *Bacillus* (99.9) |
| Female 2 | B14 | 1418 | Firmicutes | *Bacillus* (99.9) |
| Female 2 | B15 | 1031 | Firmicutes | *Bacillus* (99.9) |
| Female 2 | B16 | 1364 | Firmicutes | *Bacillus* (97.3) |
| Female 3 | B17 | 1367 | Firmicutes | *Bacillus* (99.6) |
| Female 3 | B18 | 1374 | Firmicutes | *Lysinibacillus* (97.8) |
| Female 3 | B19 | 1369 | Firmicutes | *Bacillus* (99.8) |
| Female 3 | B20 | 937 | Firmicutes | *Bacillus* (99.7) |
| Female 3 | B21 | 1366 | Firmicutes | *Bacillus* (99.9) |
| Female 3 | B22 | 912 | Firmicutes | *Bacillus* (99.7) |
| Female 4 | B23 | 1417 | Firmicutes | *Bacillus* (99.8) |
| Female 4 | B24 | 1456 | Firmicutes | *Paenibacillus* (99.7) |
| Female 4 | B25 | 1445 | Firmicutes | *Bacillus* (100) |
| Female 4 | B26 | 922 | Firmicutes | *Staphylococcus* (99.8) |
| Female 4 | B27 | 1359 | Firmicutes | *Bacillus* (99.6) |
| Female 5 | B28 | 954 | Firmicutes | *Bacillus* (99.4) |
| Female 5 | B29 | 1356 | Firmicutes | *Bacillus* (99.7) |
| Female 5 | B30 | 814 | Firmicutes | *Paenibacillus* (97.8) |
| Female 5 | B31 | 1358 | Firmicutes | *Bacillus* (99.6) |
| Female 5 | B32 | 1411 | Firmicutes | *Bacillus* (99.8) |
| Female 5 | B33 | 879 | Bacteroidetes | *Elizabethkingia* (97.8) |
| Female 5 | B34 | 1334 | Firmicutes | *Bacillus* (99.8) |
| Control 1 | B35 | 1310 | Firmicutes | *Paenibacillus* (99.3) |
| Control 2 | B36 | 804 | Firmicutes | *Bacillus* (99.6) |

**Supplementary material 3.** Dunn test comparing alpha diversity between treatments. *P* values adjusted with the Benjamini-Hochberg method are shown.

|  | Treatment 2 | Treatment 3 | Treatment 4 | Treatment 5 |
| --- | --- | --- | --- | --- |
| Treatment 1 | 0.2563 | 0.3565 | 0.4090 | 0.0209 |
| Treatment 2 | ˗̶ | 0.4074 | 0.3045 | 0.0021 |
| Treatment 3 | ˗̶ | ˗̶ | 0.4088 | 0.0077 |
| Treatment 4 | ˗̶ | ˗̶ | ˗̶ | 0.0145 |

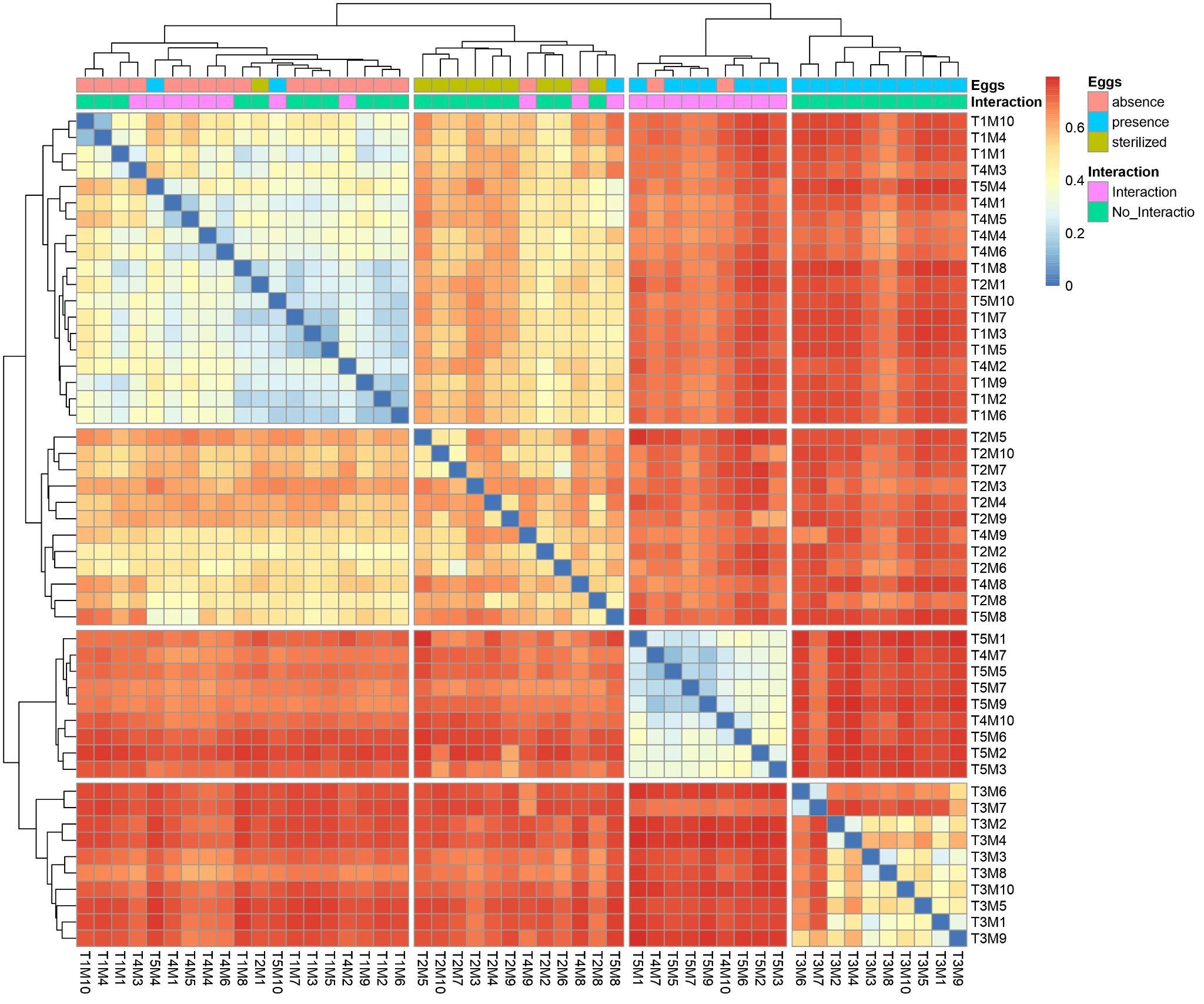

**Supplementary material 4.** Heatmap of the Jensen-Shannon distances between the five treatments.

**Supplementary material 5.** Pairwise PERMANOVA showing between-group shifts in bacterial signature profile when comparing the beta diversity among treatments. *P* values for each comparison are shown.

|  | Treatment 2 | Treatment 3 | Treatment 4 | Treatment 5 |
| --- | --- | --- | --- | --- |
| Treatment 1 | 0.001 | 0.001 | 0.001 | 0.001 |
| Treatment 2 | ˗̶ | 0.001 | 0.001 | 0.001 |
| Treatment 3 | ˗̶ | ˗̶ | 0.001 | 0.001 |
| Treatment 4 | ˗̶ | ˗̶ | ˗̶ | 0.004 |

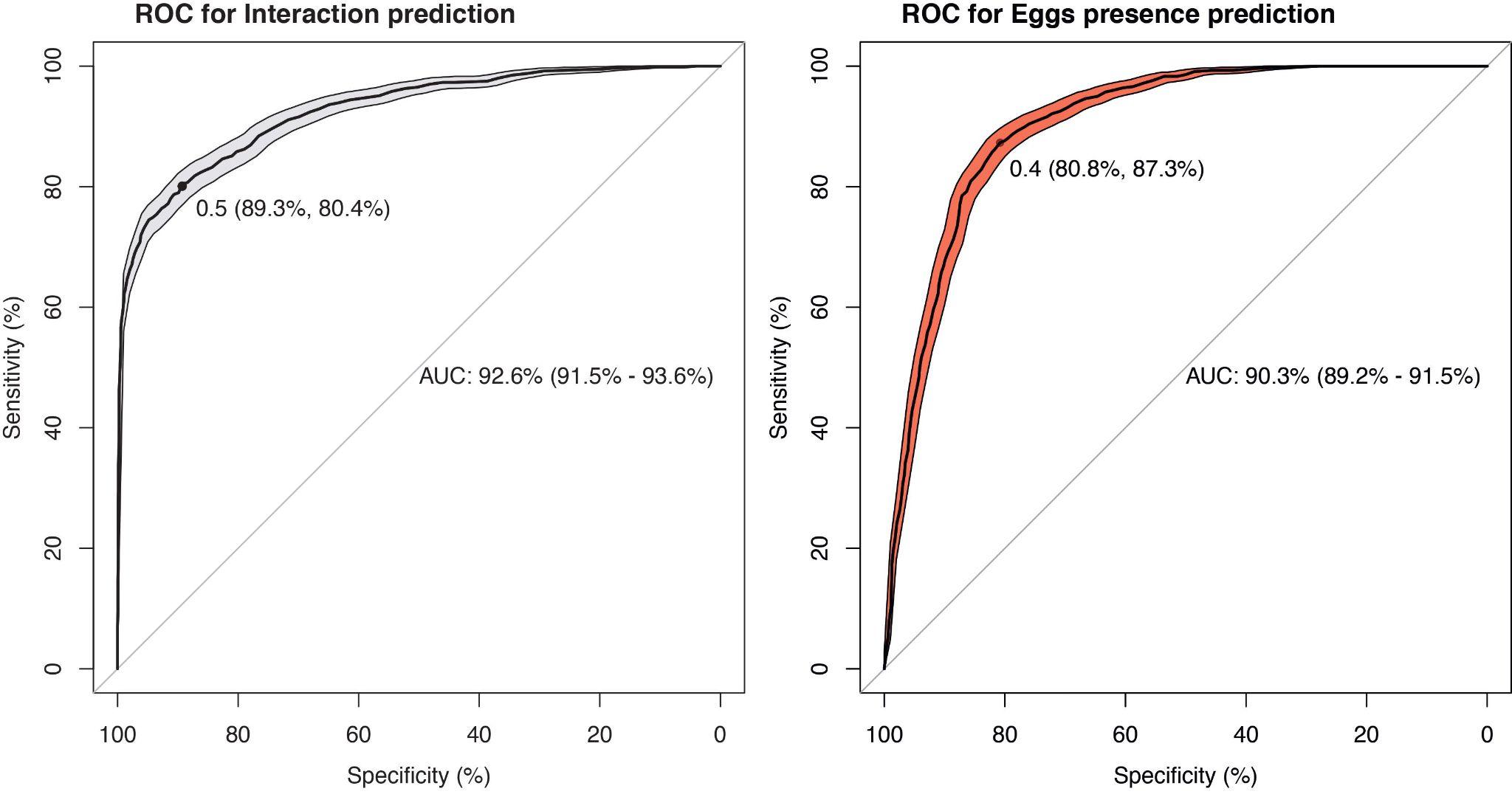

**Supplementary material 6.** ROC curves for Interaction (A) and Eggs presence (B) prediction. The point indicates the best cutoff value from the prediction probability to optimize Sensitivity and Specificity. The confidence levels reflect a 95% confidence interval.

**Supplementary material 7.** Median, mean and standard deviation (SD) of instar duration (hours) of immature stages of *Ae. aegypti* (Paea strain), wing size, and survival (95% confidence interval) exposed to *Asaia* and *Elizabethkingia* bacteria.

|  | **Control** | | | **+*Asaia*** | | | **+*Elizabethkingia*** | | |
| --- | --- | --- | --- | --- | --- | --- | --- | --- | --- |
|  | **Median** | **Mean** | **SD** | **Median** | **Mean** | **SD** | **Median** | **Mean** | **SD** |
| **L_1_ (h)** | 33 | 40.28 | 7.60 | 33 | 37.11 | 9.13 | 33 | 32.71 | 5.27 |
| **L_2_(h)** | 19 | 15.77 | 7.05 | 19 | 18.63 | 7.57 | 19 | 18.03 | 3.14 |
| **L_3_ (h)** | 24 | 24.32 | 3.55 | 24 | 24.30 | 4.89 | 24 | 23.73 | 1.46 |
| **L_4_ (h)** | 48 | 50.11 | 7.88 | 48 | 54.11 | 28.03 | 48 | 47.30 | 5.26 |
| **Pupa (h)** | 50 | 52.37 | 5.61 | 48 | 46.84 | 6.51 | 48 | 49.43 | 7.08 |
| **Total (h)** | 177 | 180.87 | 10.64 | 168 | 173.23 | 11.10 | 168 | 171.04 | 3.91 |
| **Wing length** | 2.2 | 2.27 | 0.25 | 2.1 | 2.27 | 0.28 | 2.1 | 2.27 | 0.31 |
| **Sex ratio (M:F)** | 1.9:1 | | | 2.1:1 | | | 1.7:1 | | |
| **Survival (95% CI)** | 0.85 (0.72 - 1.0) | | | 0.92 (0.84 - 1) | | | 0.77 (0.62 - 0.95) | | |

**Supplementary material 8.** Log-rank comparing the duration of all life stages.

|  |  | **Global** | **+*Asaia* vs. control** | **+*Elizabethkingia* vs. control** |
| --- | --- | --- | --- | --- |
| **L_1_-L_2_** | **Chisq** | 12.9 | 2.9 | 11.8 |
|  | **p-value** | < 0.01 | 0.08 | < 0.01 |
| **L_2_-L_3_** | **Chisq** | 0.5 | - | - |
|  | **p-value** | 0.79 | - | - |
| **L_3_-L_4_** | **Chisq** | 1.9 | - | - |
|  | **p-value** | 0.38 | - | - |
| **L_4_-Pupae** | **Chisq** | 0.9 | - | - |
|  | **p-value** | 0.65 | - | - |
| **Pupae-Adult** | **Chisq** | 7.9 | - | - |
|  | **p-value** | 0.02 | - | - |
| **L_1_-Adult** | **Chisq** | 10 | 3.7 | 12.5 |
|  | **p-value** | < 0.01 | 0.05 | < 0.01 |
| **Survival** | **Chisq** | 3.2 | - | - |
|  | **p-value** | 0.21 | - | - |

**Supplementary material 9.** Bacteria isolation from experimental groups.

| **Taxonomic identification** | **Source** |
| --- | --- |
| *Acinetobacter* | Control |
| *Bacillus* | Control, +*Asaia*, +*Elizabethkingia* |
| *Chryseobacterium* | Control |
| *Elizabethkingia* | +*Elizabethkingia* |
| *Enterobacter* | Control, +*Elizabethkingia* |
| *Exiguobacterium* | +*Elizabethkingia* |
| *Pseudomonas* | + *Asaia* |
| *Stenotrophomonas* | + *Asaia* |
